# JNK-mediated spindle reorientation in stem cells promotes dysplasia in the aging intestine

**DOI:** 10.1101/365981

**Authors:** Daniel Jun-Kit Hu, Heinrich Jasper

## Abstract

Homeostasis in high-turnover tissues depends on precise yet plastic regulation of stem cell daughter fates. In *Drosophila*, intestinal stem cells (ISCs) respond to unknown signals to switch from asymmetric to symmetric divisions during feeding-induced growth. Here, we show that this switch is controlled by dynamic reorientation of mitotic spindles by a Jun-N-terminal Kinase (JNK) / Wdr62 / Kif1a interaction. JNK promotes Wdr62 localization at the spindle and represses transcription of the kinesin Kif1a. This activity of JNK results in over-abundance of symmetric divisions in stress conditions, and contributes to the loss of tissue homeostasis in the aging animal. Restoring normal ISC spindle orientation by perturbing the JNK/Wdr62/Kif1a axis is sufficient to improve intestinal physiology and extend lifespan. Our findings reveal a critical role for the dynamic control of SC spindle orientation in epithelial maintenance.

## Introduction

SCs are critical for the regenerative capacity of many tissues, and their population size as well as their proliferative capacity has to be regulated to ensure effective regenerative responses to damage, without causing ectopic growth. Processes that ensure expansion or restoration of SC populations have been described in a variety of tissues, and may be evolutionarily conserved (O’Brien et al., 2011; Tata et al., 2013; Yan et al., 2017). Processes that increase SC numbers include de-differentiation of differentiated cell types into tissue SCs (Lucchetta and Ohlstein, 2017; Tata et al., 2013; Tetteh et al., 2016; Tian et al., 2011), as well as changes in SC division modes to ensure a higher proportion of symmetric divisions (O’Brien et al., 2011). The molecular mechanisms engaged to achieve this plasticity remain largely unclear.

Intestinal stem cell (ISC) divisions in *Drosophila* represent a productive model system to study the plasticity of SC division modes during regeneration, growth, damage, and aging. The *Drosophila* intestine is lined by a pseudo-stratified epithelium that is regenerated by ISCs after damage to ensure maintenance of barrier and digestive function (Buchon et al., 2009; Micchelli and Perrimon, 2006; Ohlstein and Spradling, 2006). ISCs contribute the vast majority of mitotic division in the intestinal epithelium, and, during asymmetric divisions, generate a new ISC and an enteroblast (EB), which terminally differentiates into one of two intestinal cell types: enteroendocrine cells (EEs) and enterocytes (ECs) (Li and Jasper, 2016; Micchelli and Perrimon, 2006; Ohlstein and Spradling, 2006). These asymmetric ISC divisions are predominant during homeostatic regeneration and after damage, but in periods of growth, the majority of ISC divisions can change to lead to symmetric outcomes. This plasticity is critical to ensure appropriate epithelial cell composition during rapid growth phases, such as after a starvation/refeeding event (O’Brien et al., 2011). The molecular and cellular mechanisms executing the switch from asymmetric to symmetric division modes have not yet been elucidated.

The cell fate of many dividing cells types, ranging from *Drosophila* neuroblasts to mammalian radial glial and epidermal basal cells, is determined by spindle orientation during mitosis (Lancaster and Knoblich, 2012; Lechler and Fuchs, 2005; Morin and Bellaïche, 2011; Siller and Doe, 2009). In dividing epithelial cells, cell polarity and spindle orientation are tightly linked, and a complex comprised of polarized cortical proteins aligns the mitotic spindle during an asymmetric division. The PDZ scaffold protein, Bazooka (Par3 in mammals), in conjunction with Par6 and atypical protein kinase C (aPKC) is enriched at the apical cell cortex, and sequesters cell fate determinants along the apical/basal axis (Bellaiche et al., 2001; Costa et al., 2008; Goulas et al., 2012; Hao et al., 2010; Wodarz et al., 1999). In *Drosophila* neuroblasts, cell fate determinants promoting differentiation versus stemness are then segregated into the two daughter cells during anaphase (Doe et al., 1991; Knoblich et al., 1995; Shen et al., 1997). To promote chromosome segregation along this axis, Par3 recruits Inscuteable (mInsc in mammals), which in turn, recruits Pins and Mud (LGN and NuMA in vertebrates)(Bowman et al., 2006; Konno et al., 2007; Postiglione et al., 2011; Siegrist and Doe, 2005; Siller et al., 2006; Williams et al., 2014). Mud, a microtubule-binding protein, interacts with the motor protein dynein to generate pulling forces along astral microtubules, reorienting the mitotic spindle towards an asymmetric division (Johnston et al., 2013; Merdes et al., 1996; Wang et al., 2011). The upstream signaling pathways that regulate the spindle orientation machinery have only recently begun to be understood.

In *Drosophila* ISCs, a role for integrins in determining spindle orientation and thus promoting asymmetric divisions has been proposed (Goulas et al., 2012). According to this model, integrin-dependent adhesion to the basement membrane results in the asymmetric segregation of Par-3, Par-6 and aPKC into the apical daughter cell, and aPKC activity then contributes to higher Notch signaling, promoting differentiation. Other studies have observed an initial symmetric segregation of Delta and Notch in the two daughter cells, and a role for transcriptional repression of Notch target genes in ISCs to preserve their identity (Bardin et al., 2010; Li and Jasper, 2016; Micchelli and Perrimon, 2006; Ohlstein and Spradling, 2006). It has further been proposed that symmetrically dividing ISCs undergo neutral competition to preserve tissue homeostasis (de Navascues et al., 2012). These models are not mutually exclusive, yet a deeper understanding of the dynamics of spindle orientation and ISC division modes is critical.

In response to stress, ISC divisions are induced by the c-Jun-N-terminal kinase (JNK) signaling pathway, both through cell-autonomous (Biteau et al., 2008; Biteau et al., 2010) and non-autonomous mechanisms (Jiang et al., 2009; Patel et al., 2015). Its activity is not required for normal regenerative proliferation, as ISCs mutant for the JNK Basket (Bsk) generate normally sized lineages under basal conditions (Biteau et al., 2008). JNK is, however, required for stress- and damage-induced proliferation (Biteau et al., 2008; Biteau et al., 2010). Chronic or excessive activation of JNK, in turn, can lead to epithelial dysplasia that is characterized by ISC over-proliferation and daughter cell mis-differentiation. This occurs during aging, where epithelial dysplasia impairs overall metabolism of the fly, disrupts intestinal barrier function, and, ultimately, shortens lifespan (Biteau et al., 2010; Rera et al., 2012). Accordingly, reducing JNK activity in ISCs extends lifespan. The mitotic response to JNK activation is mediated by the transcription factor Fos (Biteau et al., 2011), yet how JNK promotes mis-differentiation remains unclear. The JNK pathway is evolutionarily conserved, and regulates a wide array of cellular functions, including mitotic entry and, as described in the mammalian brain, spindle orientation through an interaction with the WD-repeat protein 62 (Wdr62)(Xu et al., 2014).

Here, we describe a critical role for JNK signaling in promoting symmetric ISC divisions by regulating spindle orientation in ISCs. In physiological conditions, we find that transient JNK activation is required for the symmetric outcome of ISC divisions in periods of intestinal growth during adaptive resizing of the gut. JNK regulates spindle orientation both directly, through recruitment of Wdr62 to the spindle, and transcriptionally by repressing the expression of Kif1a (Unc-104), a kinesin implicated in promoting asymmetric divisions in rat radial glial cells (Carabalona et al., 2016). We further show that, in response to excessive stress, as well as in the aging animal, chronic activation of JNK results in cell fate defects by reorienting ISC spindles planar to the basal surface. Restoring normal spindle orientation in aging and stress conditions by perturbing the JNK/Wdr62/Kif1a axis not only restores normal cell fates, but also preserves the barrier function of the intestine. Furthermore, Kif1a over-expression in ISCs also increases overall longevity. Our results deepen our understanding of cell fate regulation in barrier epithelia, and identify new intervention strategies towards preserving long-term tissue homeostasis.

## Results

### Dynamic spindle orientation in intestinal stem cells

To gain deeper insight into the cellular mechanisms controlling ISC division modes, we asked whether the switch from asymmetric to symmetric divisions during adaptive resizing (O’Brien et al., 2011) (Figure 1A) would be reflected in the spindle orientation of ISCs during mitosis. We fasted adult flies for a week, followed by refeeding to stimulate intestinal growth (Figure 1B). The low-level, homeostatic mitotic activity in the intestine of normally fed flies (quantified by phospho-histone H3, PH3, staining) declined further after 7 days of protein deprivation on 1% sucrose (fasting; Figure 1C). A 3-day refeeding period (short-term refeeding) restored mitotic activity. Consistent with this fluctuation in mitotic activity, intestinal size decreased after fasting, but returned to normal after refeeding for 7 days (long-term refeeding; Figure 1D).

**Figure 1:**
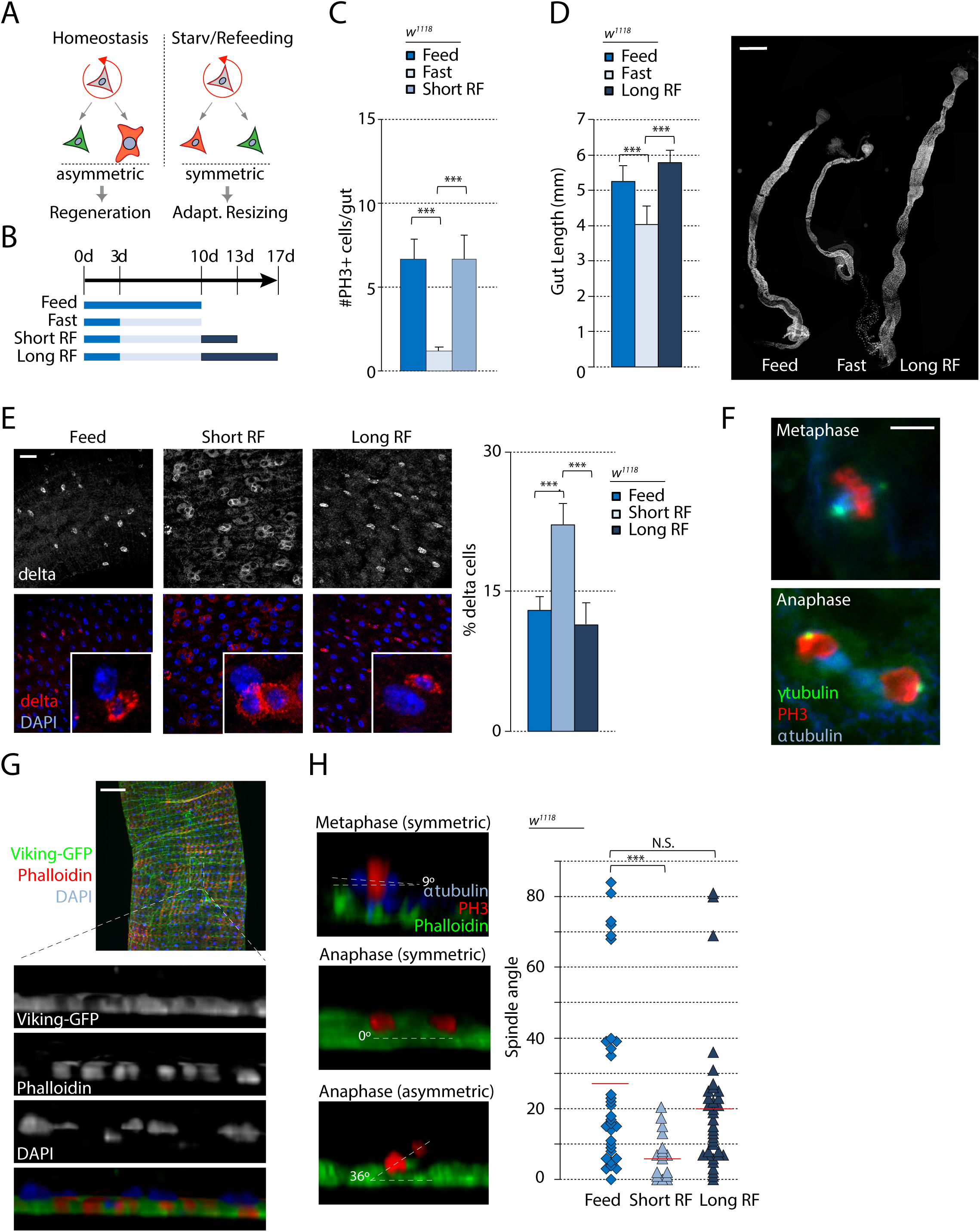
Spindle orientation is planar during adaptive intestinal growth. A. Diagram representing different modes of ISC division in response to different environmental challenges. B. Timeline of fasting and refeeding conditions. C. The number of anti-phosphohistone H3+ (PH3) cells in the entire gut decreases after fasting, but recovers after refeeding. D. *Drosophila* intestines substantially decrease in size after fasting, but recover after refeeding. E. The percentage of Delta+ cells (ISCs)/total cells in the posterior midgut increases after short-term refeeding, compared to fed flies. The percentage returns to baseline levels after long-term refeeding. Insets depict a likely clone, with two ISCs generated in short-term refeeding conditions, and an ISC and an EB generated in feeding or long-term refeeding conditions. F. Triple staining with anti-γ-tubulin, anti-PH3, and, anti-α-tubulin showing centrosomes and the microtubule network of the mitotic spindle. G. 3-D reconstruction of the intestine to visualize spindle orientation along the apical/basal axis. Fluorescently labeled collagen (Vkg) or F-actin (Phallodin staining) labels the basement membrane. H. Spindle orientations were quantified in anaphase ISCs in the posterior midgut by measuring the angle of a line bisecting segregating chromosomes with the basement membrane. Fed and long-term refed conditions favored oblique spindles while short-term refeeding favored planar spindles. mean ± SD (D,E) or mean ± SEM (C); n≥20 flies (C), n≥10 flies (D,E); N.S. = not significant, ***P<0.001, based on one-way ANOVA with Tukey test. Red bar = mean (H). Scale bar = 500μm (D), 20μm (E), 5μm (F), 50 μm (G).

Changes in intestinal size were accompanied by an increase in the percentage of Delta (Dl) expressing ISCs after 3 days of refeeding, and a return to normal levels after 7 days (Figure 1E). We asked whether this transient feeding-induced increase in Dl+ cell numbers is a consequence of symmetric divisions and is accompanied by changes in spindle orientation. We performed immunostaining against α- and γ- tubulin (microtubules and centrosomes, respectively), and quantified spindle orientation by measuring the angle between the line bisecting the two segregating sets of chromosomes during anaphase (as determined by PH3 staining) with the basement membrane (Figure 1F-1H). The orientation of the epithelial basement membrane was visualized either using Viking-GFP (a protein-trap line reporting expression of the *Drosophila* collagen Viking)(Buszczak et al., 2007) or using fluorescently-tagged Phalloidin (which visualizes the visceral muscle underlying the basement membrane; Figure 1G-1H). Mitotic ISCs in homeostatic conditions displayed oblique and widely distributed spindle angles, suggesting largely asymmetric divisions (Figure 1H). After short-term refeeding, however, most ISC spindles were grossly planar to the basement membrane (0°), suggesting a possible shift towards symmetric divisions, consistent with the observed increase in Dl+ cells. After long-term refeeding, in turn, spindles returned to largely oblique orientations, suggesting that spindle orientation in mitotic ISCs dynamically responds to growth cues, potentially controlling the fate of ISC daughter cells.

We asked whether such dynamic ISC spindle reorientation would also be observed during damage-induced regeneration, and infected flies with *Erwinia carotovora carotovora 15* (*Ecc15*), a gram-negative bacterium that damages differentiated enterocytes and induces a rapid but controlled regenerative response in the *Drosophila* intestine, before being cleared from the gut (Ayyaz and Jasper, 2013; Buchon et al., 2009). *Ecc15* exposure increased mitotic activity (Figure 2A), but did not shift spindle orientation substantially from homeostatic conditions (Figure 2B), suggesting that oblique spindles are preferred for conditions in which predominantly asymmetric ISC divisions are necessary.

**Figure 2:**
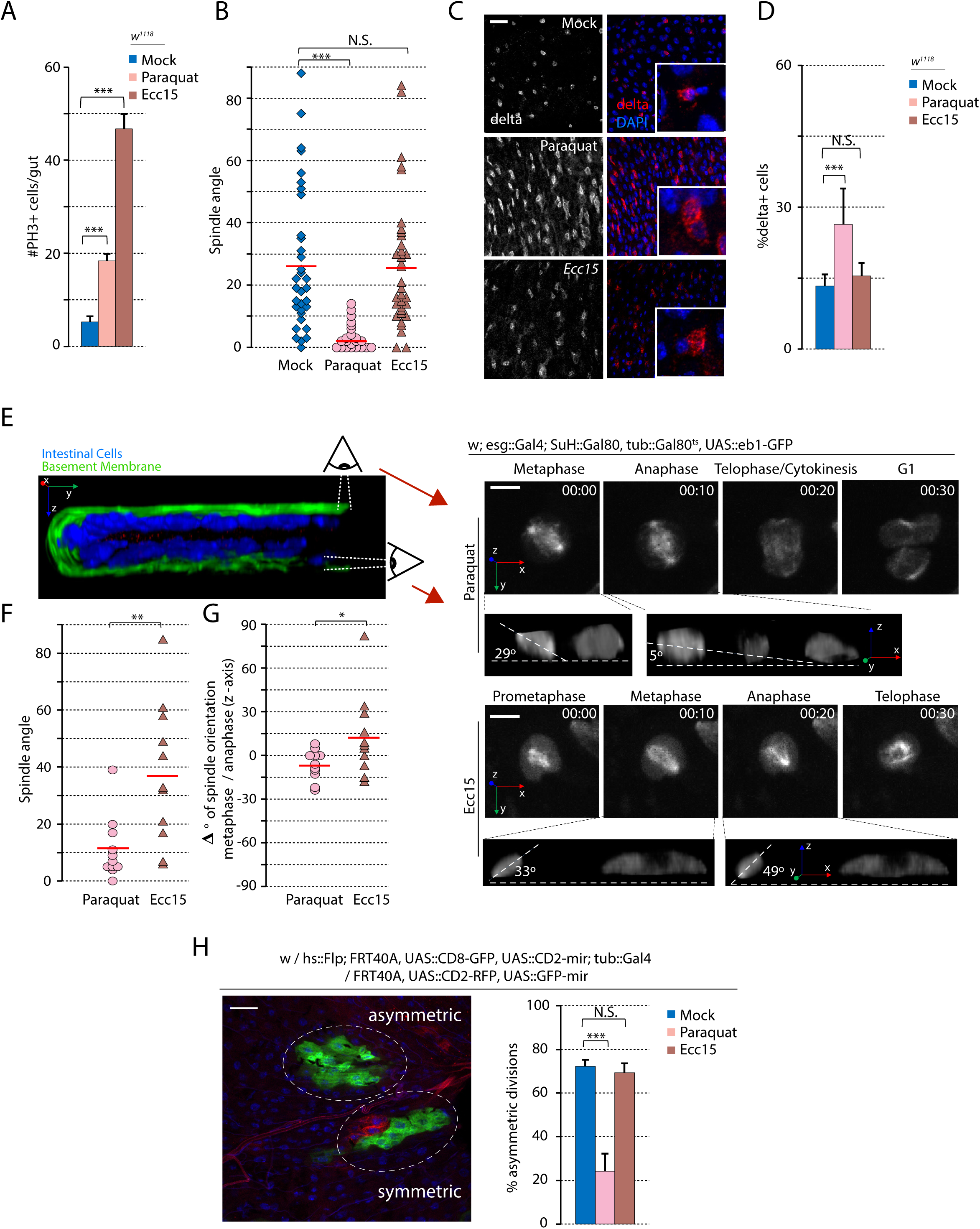
Paraquat treatment and *Ecc15* infection influences spindle orientation differently. 10d female flies were either feed with 5mM Paraquat in 5% sucrose, *Ecc15* in 5% sucrose, or 5% sucrose alone (mock control) for 20 hours. Quantifications were only performed in the posterior midgut, except for the quantification of the number of PH3+ cells. Spindle orientation was quantified during anaphase. A. Mitotic activity of ISCs in the entire gut, as determined by anti-PH3 staining, increased after Paraquat treatment or *Ecc15* infection. B. Mitotic spindles shifted to planar orientations after Paraquat treatment, but not *Ecc15* infection. C and D. The percentage of Delta+ cells (ISCs) increased after Paraquat treatment, but not after *Ecc15* infection. Insets depict a likely clone, with two ISCs generated in Paraquat-treated conditions, and an ISC and an EB generated in mock-treated or Ecc15-infected conditions. E. Intestines were dissected from flies and live imaged *ex vivo* for a 2 hour period, time (hours:minutes). Insets depicting 3D-reconstruction of cells in metaphase and anaphase to enable measurement of spindle orientation. F. Spindle orientation in live ISCs mimicked fixed cells, with largely planar spindles after Paraquat treatment and largely oblique spindles after *Ecc15* infection. G. Changes in spindle orientation between metaphase and anaphase occurred less frequently and dramatically after Paraquat treatment compared to *Ecc15* infection. In Ecc15-infected flies, the mitotic spindle of ISCs frequently oriented more perpendicular after the onset of anaphase. H. Example image of Twinspot clones to identify asymmetric (clone of only green cells) versus symmetric (clone of green and red cells) divisions. The percentage of symmetric divisions increased after Paraquat treatment, but remained unchanged after *Ecc15* infection. mean ± SD (D,H) or mean ± SEM (A); n≥20 flies (A), n≥10 flies (D,H); N.S. = not significant, *P<0.05, **P<0.01, ***P<0.001, based on one-way ANOVA with Tukey test (A,B,D,H), Student’s t-test (F,G). Red bar = mean (B,F,G,). Scale bar = 20μm (C, H), 5μm (E). See also Movies S1 and S2.

Interestingly, however, we observed a significant shift in ISC spindle orientation when subjecting flies to more severe stress conditions. Oxidative stress triggered by exposure to Paraquat induces a well-described regenerative response that is long lasting and results in dysplastic epithelial phenotypes that are reminiscent of age-related epithelial dysfunction in flies (Biteau et al., 2008). In these conditions, ISC proliferation was also increased, but spindles re-oriented planar relative to the basement membrane (Figure 2A-2B). Consistent with a symmetric outcome of these divisions, Paraquat exposure also led to an increase in relative numbers of Dl+ cells (Figure 2C-2D).

While our observations were consistent with a context-dependent reorientation of mitotic spindles that determined symmetric or asymmetric outcomes of ISC divisions, it remained unclear whether this reorientation is a consequence of modulation of fixed, pre-determining cues, or whether mitotic spindles of ISCs are oriented dynamically during mitosis. To address this question, we sought to visualize spindle orientation dynamics in stressed and infected conditions directly. Using an *ex vivo* culture system for the fly intestine (Deng et al., 2015), we used spinning-disc confocal microscopy to image mitotic events in live intestines of flies expressing a GFP-tagged version of the microtubule plus-end tracking protein EB1 in ISCs (Figure 2E-2G)(Rogers et al., 2002). EB1-GFP labeled the mitotic spindle throughout M phase, allowing dynamic tracking of spindle orientations (Figure 2E and Movie S1A-S1B). During regeneration in response to Ecc15 exposure, the spindle often rotated between metaphase and anaphase, suggesting dynamic realignment of the spindle during mitosis (Figure 2E-2G). This rotation was less common and dramatic after Paraquat treatment, and this lack of movement resulted in a more planar spindle orientation during anaphase, consistent with our observations in fixed tissue.

To confirm that the observed changes in spindle orientation are indeed instructing fate switches in the ISC lineage, we performed twinspot mosaic analysis (Yu et al., 2009). In this approach, the two daughter cells of an ISC division are differentially labeled with GFP or RFP. Subsequent lineage tracing of each daughter cell then allows to retroactively specify the fate of the two cells (Figure 2H). ISCs from flies exposed to either mock treatment or *Ecc15* infection exhibited largely asymmetric divisions (resulting in lineages in which all cells were labeled by the same color, as the differently labeled daughter cell differentiates and is eventually shed without generating a labeled clone of cells), confirming the asymmetric outcome of divisions with oblique spindle orientations (Fig. 2H). Paraquat treatment, in turn, resulted in largely symmetric divisions as represented by closely associated cell clones labeled in different colors (Figure 2H).

Altogether, these data support the idea that a productive regenerative response in the fly intestinal epithelium relies largely on asymmetric ISC divisions promoted by an actively enforced oblique spindle orientation, while in conditions in which the ISC population needs to be expanded, such as during adaptive resizing, symmetric ISC divisions are predominant due to maintenance of planar spindle orientations. Excessive stress seems to be hijacking this physiological response, preventing the acquisition of oblique orientations and thus resulting in aberrant symmetric divisions. Since such aberrant symmetric divisions are likely contributing to the dysplasia observed in aging intestinal epithelia that is recognized as a driver of mortality (Biteau et al., 2010; Rera et al., 2012), we next explored the mechanism(s) of this dynamic spindle reorientation in ISC divisions in detail.

### Activation of the JNK pathway promotes planar spindle pole orientation

The increased proliferation of ISCs in response to Paraquat depends on the activation of Jun-N-terminal Kinase (JNK) signaling, and excessive JNK activity also contributes to the elevation of Dl+ cell numbers under stress conditions (Biteau et al., 2008). Since Paraquat treatment and refeeding after starvation both resulted in planar spindles and increased frequency of symmetric divisions, we tested whether JNK may also promote symmetric ISC divisions during adaptive resizing. We assessed the presence of phosphorylated JNK (pJNK) in mitotic ISCs under varying conditions using immunohistochemistry, and found that pJNK could not be detected in mitotic ISCs under homeostasis (in normally fed animals), but was detected specifically at the spindle of ISCs after fasting/short-term refeeding (Figure 3A). This signal was not observed anymore after long-term refeeding. pJNK was also detected at the majority of mitotic ISC spindles during metaphase and anaphase in Paraquat-treated animals, but not in mock-treated or Ecc15-infected animals (Figure 3A).

**Figure 3:**
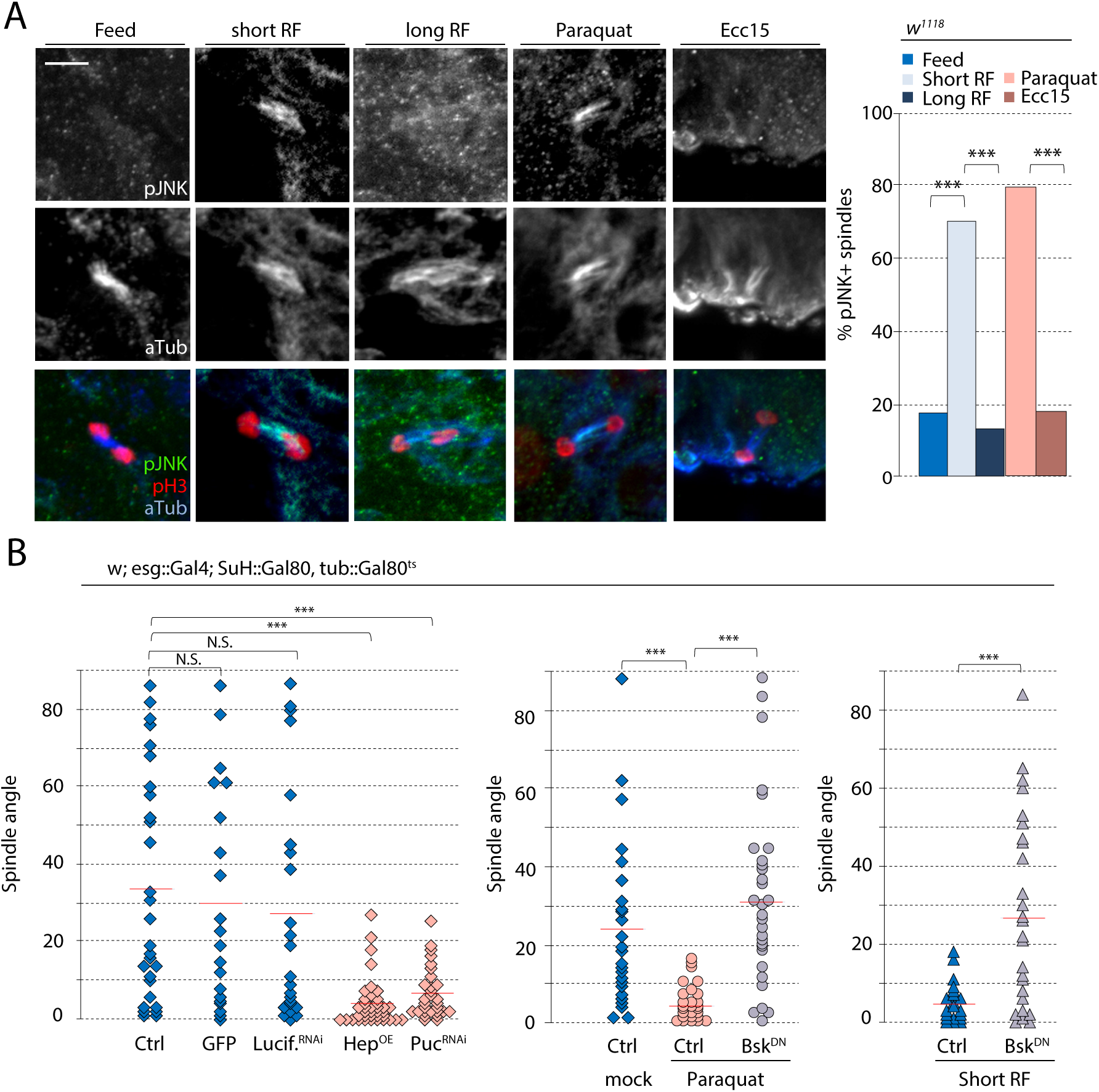
JNK activity promotes planar spindle orientation. Intestines were dissected from 10d females (except for short-term refed, Figure 1B). Only the posterior midgut was analyzed, and spindle orientation was quantified during anaphase. A. Triple staining with anti-phospho-JNK, anti-PH3, and, anti-α-tubulin revealing pJNK localization at the mitotic spindle under different environmental challenges. pJNK is normally absent at the mitotic spindle in unchallenged, fed conditions. pJNK localized to the spindle after short-term refeeding following fasting, but not after long-term refeeding. Paraquat treatment promoted pJNK localization to the spindle, but *Ecc15* infection did not. Quantification of pJNK localization at the spindle was pooled from ISCs in metaphase and anaphase. B. Overactivation of JNK by Hep^OE^ or puc^RNAi^ was sufficient to drive planar spindle orientations. Overexpression of GFP alone or Luciferase^RNAi^ had no effect on spindle orientation. Overexpression of a kinase-dead version of JNK, Bsk^DN^, was sufficient to prevent the shift to planar spindle orientation normally observed after Paraquat-treatment or short-term refeeding. n≥25 cells (A); N.S. = not significant, *P<0.05, **P<0.01, ***P<0.001, based on Chi-Squared test (A), one-way ANOVA with Tukey test (B). Red bar = mean (B). Scale bar = 5μm.

Since this recruitment of pJNK to the spindle correlated with the reorientation of spindles under the various conditions, we asked whether JNK activity controlled spindle orientation. We activated JNK in ISCs either by over-expressing a JNK kinase (*Drosophila* Hemipterous, Hep), or by knocking down *puckered* (*puc*), which encodes a JNK phosphatase (Martin-Blanco et al., 1998; Wang et al., 2003). We expressed Hep or dsRNA against *puc* inducibly and specifically in ISCs using the ISC/EB driver *escargot::Gal4* in combination with EB-specific expression of the Gal4 inhibitor Gal80 (Su(H)::Gal80) (Wang et al., 2014), and in combination with ubiquitous expression of temperature-sensitive Gal80 (tub::Gal80^ts^)(McGuire et al., 2004). Over-activating JNK specifically in ISCs increased ISC proliferation rates as previously reported (Biteau et al., 2008), and shifted spindle poles towards planar orientations, phenocopying the growth and stress conditions (Figure 3B). Conversely, disrupting the JNK pathway in ISCs by expressing a kinase-dead, dominant-negative form of the *Drosophila* JNK Basket (Bsk^DN^)(Jasper et al., 2001), limited ISC proliferation rates, and, in the remaining mitoses, also prevented the shift to planar spindle orientation after Paraquat treatment (Figure 3B). Strikingly, expression of Bsk^DN^ also prevented the planar orientation of spindles during short-term refeeding, confirming that JNK activation is critical in the switch towards symmetric divisions during adaptive resizing. Expression of Luciferase dsRNA or cytoplasmic GFP alone did not affect spindle orientation (Figure 3B). Altogether, our results suggest that the association of phosphorylated JNK with the spindle is critical for the reorientation of mitotic spindles in ISCs during stress conditions and during adaptive resizing.

### JNK biases symmetric divisions by decreasing Kif1a-mediated asymmetric divisions

JNK regulates cell function in many cases by phosphorylating transcription factors and changing the cellular transcriptome. We asked whether JNK-dependent changes in spindle orientation may also be influenced in part by the transcriptional induction or repression of selected target genes. We previously explored the transcriptome of isolated ISCs from intestines exposed to tunicamycin, which increases endoplasmic reticulum stress and triggers JNK activation in ISCs (Wang et al., 2015; Wang et al., 2014). The transcriptome changes observed in these cells compared to control conditions may thus reflect an ISC-specific JNK-induced transcriptional program. While the expression of scaffolding proteins involved in recruiting the spindle (Baz, Insc, Pins, and Mud) during asymmetric divisions was largely unaffected, Kif1a, a kinesin recently reported to promote asymmetric divisions in embryonic neural stem cells (Carabalona et al., 2016), was downregulated over four-fold (Wang et al., 2015; Wang et al., 2014). Confirming this, qPCR on RNA extracted from FACS-sorted ISCs revealed a 6-fold decrease in Kif1a RNA levels after increasing JNK activity by knocking down *puc* (Figure 4A). To assess whether this repression of Kif1a would contribute to the JNK-induced changes in spindle orientation, we tested the effect of Kif1a knockdown on spindle orientation of ISCs. Depleting Kif1a strongly increases planar spindle orientation while over-expressing Kif1a rescues the shift in planar spindle orientation caused by Paraquat or *puc* depletion (Figure 4B and 4D). These data suggest that Kif1a repression is required downstream of JNK activation to promote planar spindle orientations. Accordingly, the percentage of ISCs in Paraquat-treated intestines was decreased after Kif1a over-expression, consistent with a shift towards asymmetric divisions after promoting oblique spindle orientations (Figure 4C). Kif1a over-expression did not increase the mitotic index (Figure S1A) and did not significantly impact spindle orientation in conditions in which JNK is not activated (Figure 4B).

**Figure 4:**
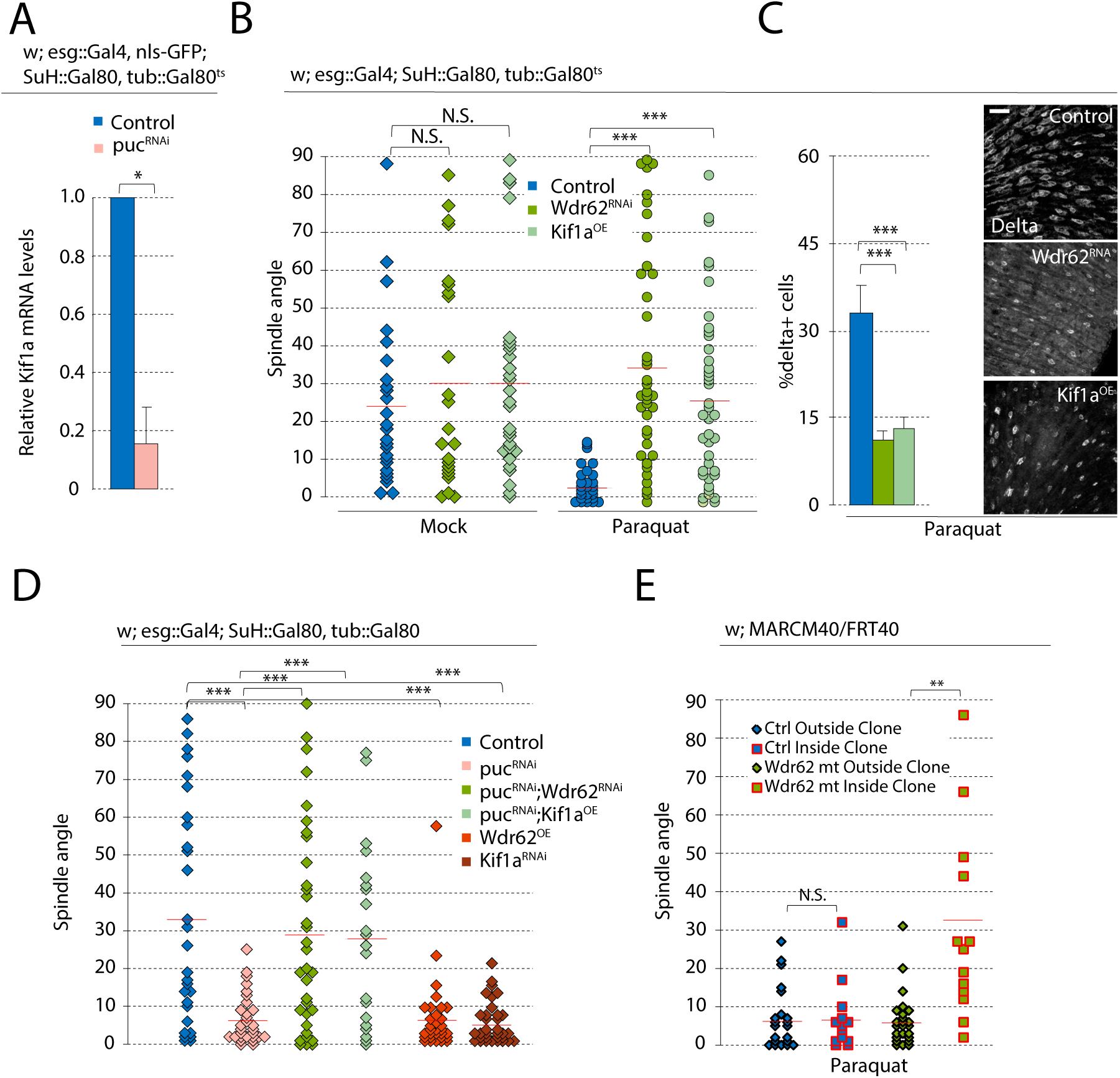
Wdr62 RNAi and Kif1a overexpression rescues stressed-induced effects on spindle orientation. Intestines were dissected from 10d old females. Only the posterior midgut was analyzed. Spindle orientation was only quantified during anaphase. A. Relative mRNA levels of Kif1a, normalized to Actin5c, in ISCs were decreased after puc^RNAi^. B. In mock-treated conditions, Wdr62^RNAi^ or Kif1a^OE^ did not affect spindle orientation. In Paraquat-treated conditions, Wdr62^RNAi^ or Kif1a^OE^ prevented the shift to planar spindle orientation. Mock-treated and Paraquat-treated controls data were taken from Figure 3B, as experiments were done in parallel. C. The percentage of Delta+ cells (ISCs) was reduced after Wdr62^RNAi^ or Kif1a^OE^ in Paraquat-treated conditions. D. Co-expression of either Wdr62^RNAi^ or full-length Kif1a with puc^RNAi^ was sufficient to rescue the shift to planar spindle orientation observed after puc^RNAi^ alone. Wdr62^OE^ or Kif1a^RNAi^ in unchallenged conditions promoted planar spindle orientations. Untreated control and puc^RNAi^ data were taken from Figure 3B. E. *wdr62* loss of function ISCs rescued the Paraquat-dependent shift to planar spindles. mean ± SEM (A) or mean ± SD (C); n≥10 flies (C); N.S. = not significant, *P<0.05, **P<0.01, ***P<0.001, based on paired, two-tailed t-test (A), one-way ANOVA with Tukey test (B,C,D), Student’s t-test (E). Red bar = mean (B,D,E). Scale bar = 20μm. See also Figure S1.

### JNK is recruited to ISC spindle by Wdr62 to promote symmetric divisions

A role for JNK signaling in modulating spindle orientation has previously been described in the developing mammalian cortex, in which JNK was shown to be recruited to the spindle of mitotic radial glial cells by the WD40-repeat protein Wdr62 (Xu et al., 2014). Wdr62 is a centrosome-associated protein and a phosphorylation target of JNK (Bogoyevitch et al., 2012; Cohen-Katsenelson et al., 2011; Williams et al., 2011; Williams et al., 2014). Loss of Wdr62 was reported to cause cell fate defects in both *Drosophila* and mammalian neural stem cells (Jayaraman et al., 2016; Lim et al., 2017; Ramdas Nair et al., 2016; Xu et al., 2014), but the role of Wdr62 on spindle orientation has yet to be examined in non-neuronal tissue. Given the observed role for JNK in regulating ISC spindle orientation, we tested whether Wdr62 would also influence spindle orientation in these cells. Indeed, over-expression of Wdr62 in ISCs shifted spindle orientation planar to the basal surface, while depleting Wdr62 by RNAi in ISCs was sufficient to prevent the JNK- or Paraquat-induced shift towards planar spindle orientation (Figure 4B and 4D). Consistent with these changes in spindle orientation, the percentage of ISCs in intestines of Paraquat-treated animals was lowered after Wdr62 depletion (Figure 4C). Depleting Wdr62 in mock-treated conditions did not significantly change average spindle pole angles (Figure 4B), suggesting that Wdr62 is specifically engaged to control spindle orientation in stress conditions that also activate JNK. As an alternative strategy to test Wdr62 function, we generated ISC lineages homozygous for a *wdr62* loss-of-function allele (wdr62^Δ3^*^−9^*)(*Ramdas* Nair et al., 2016) using MARCM (mosaic analysis with a repressible cell marker) (Lee and Luo, 2001), and compared spindle orientation between mitotic cells within clones (i.e. Wdr62 deficient ISCs) and outside clones (i.e. wild-type ISCs). After Paraquat treatment, spindle orientation from control crosses were largely planar in mitotic cells both inside and outside clones (Figure 4E). In MARCM clones homozygous for *wdr62*^Δ^*^3-9^*, however, spindle orientation was largely oblique even after Paraquat exposure, while spindle orientation of mitotic ISCs outside of clones in flies of the same genotype shifted to planar orientations as expected. In mouse embryonic neural stem cells, loss of Wdr62 has been reported to cause defects in spindle assembly in addition to changes in spindle orientation, though its exact role remains controversial (Jayaraman et al., 2016). In the *Drosophila* intestine, spindle assembly was not obviously impacted after Wdr62 knockdown (Figure S1B), nor were numbers of mitotic cells different compared to controls (Figure S1A), suggesting that, in ISCs, Wdr62 plays a specific role in the orientation of the mitotic spindle. Clone sizes from *wdr62* mutant ISCs were reduced compared to controls (Figure S1C), yet it remains unclear whether this is caused by defects in mitotic progression or due to decreased symmetric divisions.

We then examined the relationship of JNK and Wdr62 at the spindle. Paraquat-mediated accumulation of pJNK at the spindle was not observed in animals expressing Bsk^DN^ or Wdr62^RNAi^ (Figure 5A), while knockdown of *puc* resulted in pJNK localization to the spindle in untreated animals. Similar to pJNK, Wdr62 was present at the spindle after Paraquat treatment, but not after *Ecc15* infection (Figure 5B), and in Paraquat-exposed animals expressing Bsk^DN^, Wdr62 was not observed at the spindle (Figure 5C). Depletion of Wdr62 itself also resulted in loss of the Wdr62 signal, confirming the specificity of the antibody (Figure 5C). Furthermore, increasing JNK activity by depletion of Puc was sufficient to promote Wdr62 localization to the spindle. Elevated JNK activity is thus sufficient to recruit Wdr62 to the mitotic ISC spindle, and phosphorylated JNK and Wdr62 are mutually dependent on each other for their translocation to the spindle under stress conditions, where they promote planar spindle orientation.

**Figure 5:**
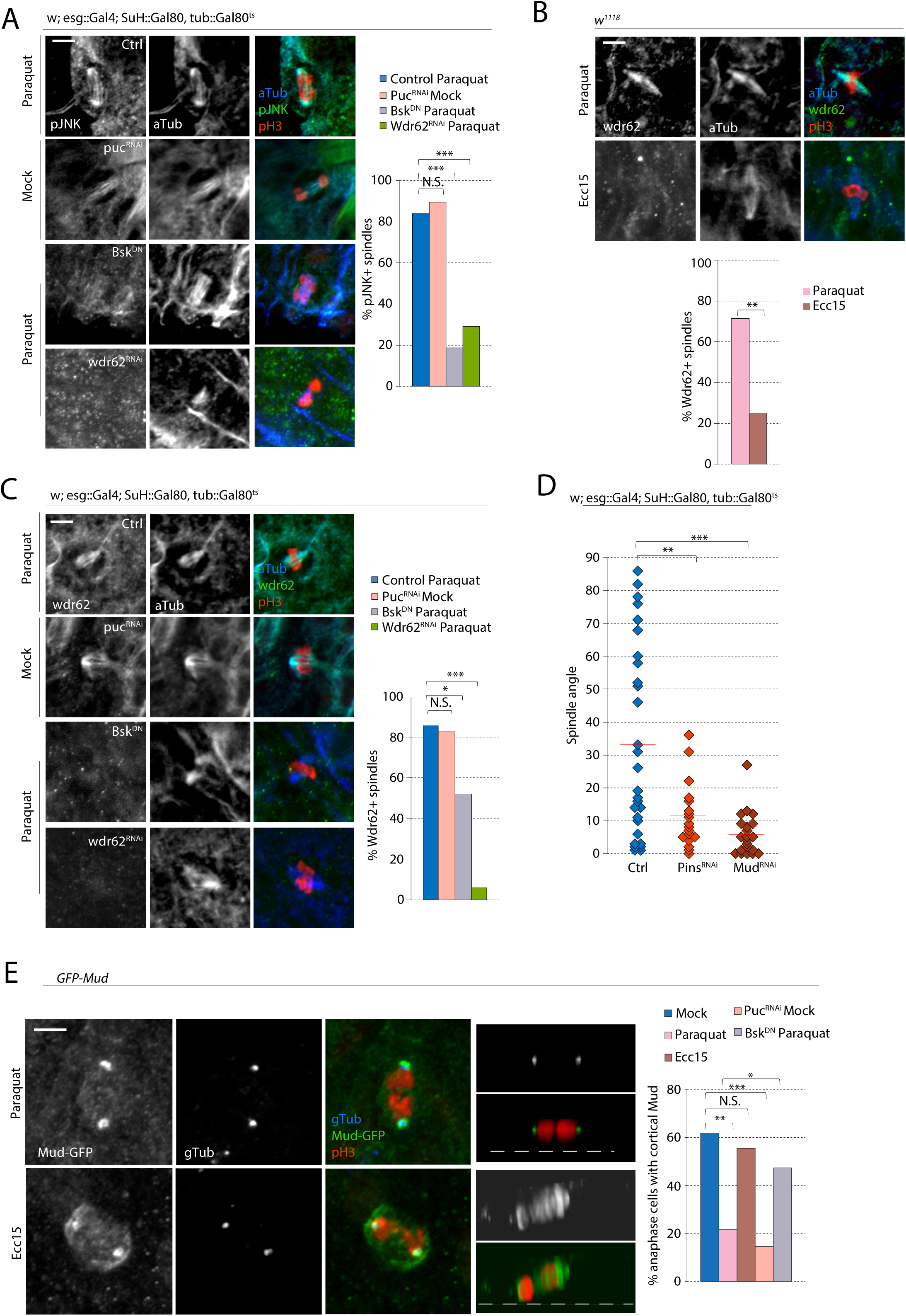
phospho-JNK and Wdr62 localization to the spindle prevents the recruitment of cortical Mud. Intestines were dissected from 10d old females. Only the posterior midgut was analyzed, and spindle orientation was only quantified during anaphase. A. JNK activity at the mitotic spindle was identified by co-localization of pJNK staining with α-tubulin staining in PH3+ cells. In control crosses, pJNK localized to the spindle after Paraquat-treatment. pJNK localization was lost after expression of BskDN or Wdr62 RNAi. puc^RNAi^ promoted pJNK localization to the ISC spindle. Quantification of pJNK localization at the spindle was pooled from ISCs in metaphase and anaphase. B. Triple staining with anti-Wdr62-JNK, anti-PH3, and, anti-α-tubulin revealed Wdr62 localization at the mitotic spindle after inducing Paraquat-treatment, but not after *Ecc15* infection. Quantification of Wdr62 localization at the spindle was pooled from ISCs in metaphase and anaphase. C. The percentage of mitotic spindles with Wdr62 localization decreased after expression of Bsk^DN^ or Wdr62^RNAi^. D. Depletion of components of spindle recruitment machinery to the apical membrane caused a loss of oblique spindles. E. Overexpression of GFP-tagged Mud revealed localization at both the centrosome and the cell cortex, or only the centrosome. After *Ecc15* infection, Mud was localized at both the cell cortex and centrosomes in the majority of anaphase cells. After Paraquat treatment, cortical Mud was absent in the vast majority of anaphase cells. puc^RNAi^ phenocopied Paraquat treatment while expression of Bsk^DN^ rescued the loss of cortical Mud from Paraquat treatment. n≥20 cells (A,B,C,E); N.S. = not significant, *P<0.05, **P<0.01, ***P<0.001, based on Chi-Squared test (A,B,C,E) and one-way ANOVA with Tukey test (D). Red bar = mean (D). Scale bar = 5μm.

### Mud is sequestered away from the cortex in stressed ISCs

To better understand how JNK regulates spindle orientation, we examined proteins involved in anchoring astral microtubules, and thus in linking the spindle with the cell cortex. We tested the role of two such proteins, Pins and Mud (LGN and NuMA in mammals) (Lancaster and Knoblich, 2012; Siller and Doe, 2009). Depleting either Pins or Mud through RNAi increased planar spindle orientation, suggesting that both are required to allow oblique spindle orientation (Figure 5D). Because Mud also plays additional roles at the spindle pole during spindle assembly (Bosveld and Ainslie, 2017; Du et al., 2002; Izumi et al., 2006; Silk et al., 2009), we overexpressed, using a Bac-based transgene GFP-tagged Mud (Bosveld et al., 2016) to visualize its localization and differentiate between the two functions of Mud. After inducing a regenerative response by *Ecc15* treatment, we found that 55% of ISCs had Mud localization at both the spindle pole and the cell cortex (Figure 5E). After Paraquat treatment, however, Mud was largely localized to the mitotic spindle, suggesting a potential role for JNK-mediated modulation of Mud localization to promote planar spindle orientation. Indeed, *puc* depletion was sufficient to cause a loss of cortical Mud in untreated conditions, while Mud was retained at the cell cortex after Paraquat treatment when Bsk^DN^ was expressed (Figure 5E). These data suggest that the activity of JNK promotes release of Mud from the ISC cell cortex to promote symmetric divisions.

### Wdr62 and Kif1a regulation is required for symmetric divisions during periods of growth

Together, these results suggest that JNK promotes symmetric division of ISCs by at least two mechanisms: a) through its direct function at the spindle where it promotes Wdr62 recruitment and modulates Mud localization, and b) through its downregulation of Kif1a expression to prevent oblique spindle orientations. We asked whether this interaction of JNK with Wdr62 and Kif1a would also be observed during adaptive resizing, a condition that engages JNK activity in a more physiological setting than Paraquat exposure. Similar to Bsk^DN^ expression (Figure 3B), knock down of Wdr62, or Kif1a over-expression, prevented the planar shift of spindle poles after fasting/short-term refeeding (Figure 6A), and resulted in a lower percentage of ISCs compared to controls (Figure 6B). Wdr62 depletion or Kif1a over-expression did not affect mitotic activity during the refeeding process (Figure 6B). We then examined the consequences of preventing symmetric divisions during growth. Because JNK activity is required for the induction of ISC mitoses, and as such, would affect tissue growth regardless of its effect on spindle orientation, we focused on Wdr62 and Kif1a. After subjecting ISCs to Wdr62 knock down or Kif1a over-expression during the refeeding process, intestines failed to return to normal size, resulting in decreased tissue length compared to control (Figure 6C).

**Figure 6:**
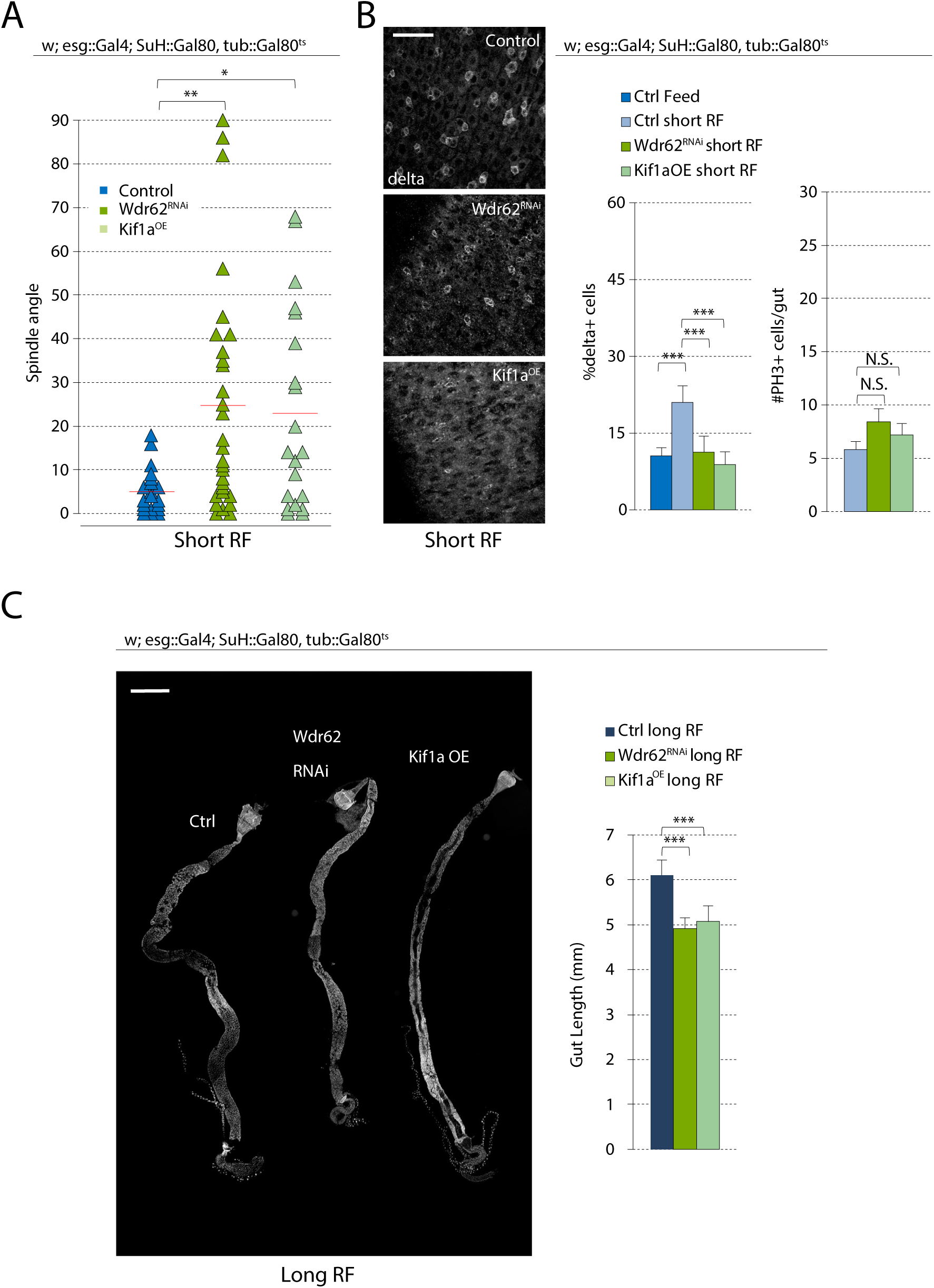
Regulation of spindle orientation during adaptive growth is dependent on Wdr62 and Kif1a. Intestines were dissected from females at ages described in Figure 1B. Only the posterior midgut was analyzed, except for PH3 counts. Spindle orientation was only quantified during anaphase. A. Wdr62^RNAi^ and expression of full-length Kif1a was sufficient to prevent the shift in planar spindle orientation observed after short-term refeeding. Control data for short-term refeeding were taken from Figure 3B, justified by the pooling of cells from a large number of flies over many experiments (over 25 flies, excluding those without anaphase cells in the posterior midgut, from 7 independent experiments). B. The percentage of Delta+ cells (ISCs) was reduced after Wdr62^RNAi^ or Kif1a overexpression in short-term refed conditions. The number of PH3+ cells in the gut was not affected. C. Gut length was decreased after Wdr62^RNAi^ or Kif1a overexpression in long-term refed conditions. mean ± SD (B: %Delta+ cells, C) or mean ± SEM (B: #PH3+ cells); n≥10 flies; N.S. = not significant, **P<0.01, ***P<0.001, based on one-way ANOVA with Tukey test. Red bar = mean (A). Scale bar = 20μm (B), 500μm (C).

### Spindle orientation in aging ISCs is chronically shifted planar to the basal surface

While various environmental challenges activate ISC mitoses, we have found that cell fate decisions in the ISC lineage are altered depending on the stimulus: symmetric divisions prevail during growth, while asymmetric divisions predominate during regeneration. We hypothesized that this regulation is crucial in maintaining tissue homeostasis. Supporting this view, when ISCs favor symmetric divisions in response to excess stress signals (as after Paraquat treatment), dysplasia is observed (Figure 2)(Biteau et al., 2008). A similar phenotype is observed in aging animals, where chronic activation of JNK as a consequence of commensal dysbiosis, epithelial inflammatory responses, and increased ER and oxidative stress, triggers increased ISC proliferation and mis-differentation of ISC daughter cells, resulting in increased numbers of Dl+ cells (Guo et al., 2014; Li et al., 2016). Since this phenotype is reminiscent of the consequences of promoting a planar spindle orientation in ISCs, we asked whether the intestinal epithelium of old animals exhibits an increase in planar oriented ISC spindles. We confirmed that mitotic activity and the percentage of Delta+ cells are increased in old animals, and found that spindle orientation in a majority of ISCs were planar in old flies, compared to the oblique orientation predominantly found in ISCs of young animals (Figure 7A). We tested whether the over-activation of JNK and its downstream effects contributed to these excessive symmetric divisions, and expressed Bsk^DN^ in ISCs using the ISC/EB-specific, RU486 inducible, 5961-Geneswitch expression system (Mathur et al., 2010). This driver allows comparing chronic effects of genetic perturbations in ISCs in genetically identical siblings. RU486 did not change the trends in spindle orientation in young versus old ISCs in wild-type backgrounds, yet after inducing Bsk^DN^ expression at 36 days of age, spindle orientation was rescued in 40 day old animals (Figure 7B). Similarly, depleting Wdr62 or overexpressing Kif1a was sufficient to rescue spindle orientation defects in old animals. Restoring spindle orientation in old animals was also sufficient to lower the abnormally high percentage of ISCs (Figure 7C). Our data, therefore, provide evidence that chronic JNK activation, and its interaction with Wdr62 and repression of Kif1A expression, cause aberrant cell fates in the aging intestinal epithelium by promoting planar spindle orientations. Furthermore, successful rescue after only short-term intervention suggests that the defects in spindle orientation are not irreversible.

**Figure 7:**
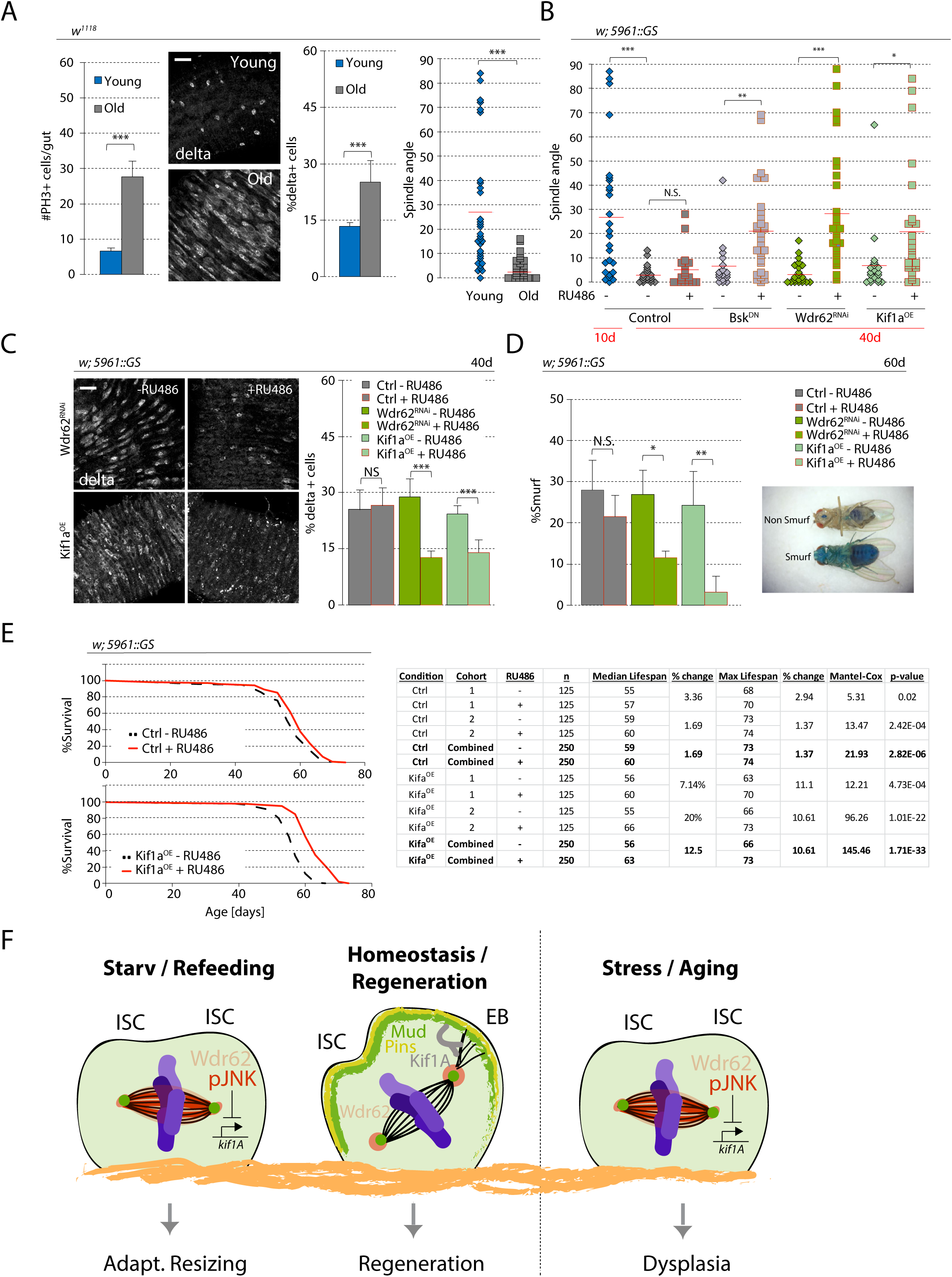
Rescuing age-associated spindle orientation defects restores intestine physiology. Intestines were dissected from either 10d (young) or 40d (old) females. When gene activity was under the 5961::GS promotor, expression was activated by 200uM RU486 in normal food (see Materials and Methods). Only the posterior midgut was analyzed, except for PH3 counts. Spindle orientation was only quantified during anaphase. A. Mitotic activity and the ratio of ISCs increased in old flies, as determined by anti-PH3 staining and anti-Delta staining respectively. Spindle orientation of mitotic ISCs was dramatically shifted planar in old flies. Spindle orientation for young flies were taken from Figure 1H (fed flies), as young flies and fed flies are the same condition, and data was pooled from a large number of flies over many experiments (18 flies, excluding those without anaphase cells in the posterior midgut, from 8 independent experiments). B. RU486 was utilized to activate gene expression in 36d old flies, and intestines were dissected from 40d old flies. Spindle orientation was widely distributed and largely oblique in young flies, but shifted to planar orientations as flies aged. RU486 alone did not affect spindle orientation. Activating Bsk^DN^, Wdr62^RNAi^, or full-length Kif1a expression with RU486 was sufficient to rescue largely oblique spindles. C. RU486 was utilized to activate gene expression in 36d old flies, and intestines were dissected from 40d old flies. RU486 alone did not affect the ratio of ISCs (as determined by anti-Delta staining) in aging flies, but activating Wdr62^RNAi^ or full-length Kif1a expression decreased the percentage of Delta+ cells in the aging intestine. D. RU486 mixed with Blue dye no.1 in normal food was fed to 36d females continuously until 60d. Intestinal barrier function was assayed by the presence of blue dye strictly in the intestine versus the entirety of the fly (a Smurf fly). 30% of flies were ‘Smurfs’ by day 60 and RU486 alone had no significantly effect. Activation of Wdr62 RNAi or full-length Kif1a expression reduced the percent of Smurf flies, extending barrier function. E. RU486 was fed to 10d females continuously until the end of the analysis. RU486 alone had minimal effect on lifespan (<2% median and maximum lifespan increase). Kif1a full-length expression resulted in substantial increase in lifespan (12.5% median and 10.6% maximum). Each graph contains data pooled from two independent experiments with details of statistical analysis provided in the chart. F. Schematic representation of JNK-mediated regulation of spindle orientation in starved/refeeding, stressed, and aging conditions through its interaction with Wdr62 to prevent recruitment of cortical Mud, and downregulation of *kif1a* transcript levels. mean ± SD (A: %Delta+ cells,C,D) or mean ± SEM (A: #PH3+ cells); n≥10 flies (A: %Delta+ cells, C), n=3 cohorts; ≥ 30 flies per cohort (D), n≥20 cells (A: %Spindle Angle); N.S. = not significant, *P<0.05, **P<0.01, ***P<0.001, based on Student’s t-test. Red bar = mean (A,B). Scale bar = 20μm. See also Figure S2.

### Restoring chronic planar spindle orientation extends tissue function and lifespan

As the fly ages, barrier function of the intestine declines, resulting in loss of intestinal integrity and imminent death (Rera et al., 2012). We tested whether maintaining proper cellular composition by manipulating cell fates through modulating spindle orientation would prevent this barrier dysfunction. To visualize the loss of barrier function, flies were fed with a non-absorbable blue dye (FD&C blue dye no. 1) starting from 30 days of age. In a healthy fly, only the intestine is stained blue, but when barrier function is lost, the entire fly turns blue, a phenotype referred to as ‘Smurf’ (Rera et al., 2011) (Figure 7D). When knock down of Wdr62 or over-expression of Kif1a was induced starting at 30 days of age, a significant decrease of Smurf flies was observed at 60 days of age (Figure 7D). RU486-treatment in control flies did not significantly affect the number of Smurf flies. Because intestinal health strongly influences *Drosophila* lifespan, we examined whether restoring spindle orientation could extend lifespan. Flies were fed RU486 to induce expression starting at 10 days of age, and while RU486 treatment in the control cross caused only minor changes in median lifespan (1.6%), Kif1a expression increased median lifespan by 12% (Figure 7E). Depletion of Wdr62, however, inconsistently affected lifespan in different cohorts (Figure S2), possibly because Wdr62 is known to play other crucial centrosomal functions during interphase (Jayaraman et al., 2016; Ramdas Nair et al., 2016). Combined, our results show that manipulating spindle orientation is sufficient to prevent the loss of tissue homeostasis, and demonstrate the feasibility of perturbing spindle orientation to extend tissue function and increase longevity.

## Discussion

A conserved role of spindle positioning in the control of asymmetric cell divisions during development and in tissue homeostasis has been established in various systems (Morin and Bellaïche, 2011; Siller and Doe, 2009), and disruption in the stereotypical positioning of the spindle in male *Drosophila* germline stem cells has been attributed to declining sperm production with age (Cheng et al., 2008). A context-dependent transition from asymmetric to symmetric divisions has further been described during adaptive resizing in the *Drosophila* intestine (O’Brien et al., 2011), yet the role of spindle repositioning in this plasticity remained unclear. Our study identifies a function for JNK signaling in promoting symmetric divisions through the realignment of the mitotic spindle during adaptive resizing and after stress, and highlights the contribution of this mechanism to the loss of tissue homeostasis in the aging intestine. Our data support a model in which the mutual recruitment of phospho-JNK and Wdr62 to the spindle, as well as the JNK-dependent transcriptional repression of Kif1a are required for spindle repositioning towards a planar orientation. Since the activation of JNK also prevents cortical localization of Mud, we propose that JNK activity disrupts engagement of the spindle with cortical determinants of spindle orientation by preventing the interaction of cortical Mud, and limits the force exerted on astral microtubules by repressing Kif1A expression (Figure 7F). Our live imaging analysis suggests that this prevents the active realignment of the spindle during the metaphase to anaphase transition, resulting in increased symmetric outcomes of ISC divisions, and thus an amplification of the stem cell population.

### Cell fate regulation by spindle orientation

The role of regulated spindle repositioning in promoting symmetric ISC divisions during adaptive resizing has parallels in developing mammalian tissues, in which spindle orientation is mostly planar early in development to increase the stem cell population before switching to mostly oblique spindles (Haydar et al., 2003; Lechler and Fuchs, 2005; Williams et al., 2014). Also in the development of the fly intestine, spindle orientation is regulated to ensure appropriate cell composition of the tissue: during pupation, in which ISCs give rise only to entero-endocrine cells, planar spindle orientations are nearly absent, even compared to the adult intestine (Guo and Ohlstein, 2015). In contrast to these developmentally choreographed changes in mitotic spindle orientation, our data indicate that spindle orientation is plastic and responds to environmental challenges in the adult intestinal epithelium. The disparity between spindle behaviors after Paraquat treatment and *Ecc15* infection shows that the nature of the environmental trigger is critical. While both stresses induce strong proliferative responses, their effects on spindle orientation and the corresponding cell fate outcome are very different.

Our live imaging results reveal that the mitotic spindle is not preset at the onset of mitosis, but is dynamically repositioned throughout metaphase and anaphase during the regenerative response to *Ecc15* infection. Paraquat treatment, which induces JNK activity in ISCs, in turn, results in a mitotic spindle that is less dynamic and favors more planar orientations. This suggests that a planar orientation may be the default during early mitosis, and that for a regenerative response that requires asymmetric divisions, the spindle is actively rotated during metaphase and anaphase to acquire an oblique angle. Because planar spindles are more dominant during periods of growth (including development), this orientation may be the physiological default state in somatic stem cell divisions. The requirement for JNK activation in accomplishing planar positioning is interesting in this context, and the identification of physiological trigger(s) of JNK activation at the metaphase-to-anaphase transition will be of significant interest in future work. Furthermore, how the dynamic repositioning of the spindle during asymmetric ISC mitoses is ensured, and whether the flexibility of spindle positioning is critical for ISC function, will be interesting questions to address.

### Role of JNK and Wdr62 at the mitotic spindle

The recruitment of phospho-JNK and Wdr62 to the spindle are mutually dependent on each other. However, the magnitude of the effect of JNK loss on the association of Wdr62 with the spindle was less compared to the effect of Wdr62 loss on pJNK localization. The more modest decrease may be indicative of a role for other kinases that have been reported to recruit Wdr62 to the centrosome, including Aurora A and Polo-like kinase (Lim et al., 2017; Lim et al., 2016; Miyamoto et al., 2017; Ramdas Nair et al., 2016). Nonetheless, JNK seems to play a role in stabilizing Wdr62 at the spindle, as JNK over-activation was sufficient to increase Wdr62 affinity with the spindle.

The role of Wdr62 in stem cell activity has previously been characterized in the developing brain (Bogoyevitch et al., 2012; Jayaraman et al., 2016; Lim et al., 2017; Nicholas et al., 2010; Ramdas Nair et al., 2016; Shohayeb et al., 2017; Xu et al., 2014; Yu et al., 2010), and our study marks the first study supporting a role of Wdr62 in spindle orientation of adult, non-neuronal stem cells. Mutations in Wdr62 are a hallmark of microcephaly, where the developing brain fails to reach its proper size because stem cell production is stunted (Bilgüvar et al., 2010; Nicholas et al., 2010; Yu et al., 2010). The fact that overexpressing Wdr62 resulted in mostly symmetric divisions (Figure 3D) suggests that Wdr62 levels must be carefully regulated to alternate between symmetric and asymmetric divisions. Unlike other reports in neural stem cells (Jayaraman et al., 2016; Ramdas Nair et al., 2016), we did not find an obvious role of Wdr62 in maintaining bipolar spindles, although it is unclear whether this is a result of different genetic tools, or a reduced role of Wdr62 in either adult, non-neuronal, or *Drosophila* tissue. Reports have also identified roles of Wdr62 in stabilizing microtubules and centrosomes in interphase neural stem cells (Jayaraman et al., 2016; Ramdas Nair et al., 2016), and while we did not explore the effects of Wdr62 depletion during interphase, the absence of gross mitotic defects suggests that in *Drosophila* ISCs, Wdr62 may function selectively in mitotic spindle orientation. On the other hand, we have observed somewhat smaller clone sizes of ISC lineages deficient for Wdr62, and therefore cannot rule out an important function for interphase Wdr62. Disruption of Wdr62 activity during interphase may also contribute towards the inconsistent effect on lifespan observed after Wdr62 depletion, despite the restoration of oblique spindles in ISCs of old flies. Further studies broadening the characterization of Wdr62 function in ISCs will be of interest.

The fact that depletion of Pins and Mud resulted in a bias towards planar spindle orientation that phenocopies the activation of JNK, and the observation that JNK activation prevents cortical Mud localization, further supports the notion that JNK actively prevents the interaction between astral microtubules and the cortical cytoskeleton to promote symmetric divisions. The consequences of the loss of Pins and Mud vary depending on the tissue: disrupting Pins and Mud in *Drosophila* neuroblasts randomizes the mitotic spindle, but, in the mammalian skin, basal stem cells with depleted LGN or NuMA favor planar spindles, similar to our observations in *Drosophila* ISCs (Siller et al., 2006; Williams et al., 2011). We propose that activated JNK and Wdr62 are recruited to the spindle in part to sequester Mud away from the cell cortex to promote planar spindles. The extent to which JNK or Wdr62 interact directly with Mud is an important question for further study.

### Role of Kif1a in asymmetric divisions

Similar to mammalian radial glial cells (Carabalona et al., 2016), Kif1a in ISCs promoted oblique spindle orientation, although the mechanism by which the kinesin does so is unclear. Khc-73, a kinesin in the same Kinesin-3 family (Hirokawa et al., 2009), is believed to interact with Pins or Disc Large in *Drosophila* S2 cells and neuroblasts to orient astral microtubules to the cell cortex, and Kif1a may play similar roles in ISCs (Johnston et al., 2009; Siegrist and Doe, 2005). While our data suggest that JNK regulates Kif1a levels transcriptionally, it is possible that JNK also regulates Kif1a at the protein level, and may direct its possible interaction with the spindle recruitment machinery. Kif1a also participates in vesicular transport in non-neuronal and neuronal cells, and may thus affect spindle orientation indirectly during interphase (Xue et al., 2010; Yonekawa et al., 1998). The role of Kif1a during interphase ISCs, if any, are unknown. Because Kif1a manipulations had a stronger effect on barrier function and longevity than Wdr62, understanding the mechanisms of how Kif1a affects spindle orientation is important to better understand the aging condition.

### JNK activity and spindle orientation during tissue resizing and aging

Our data highlight how a physiologically important role for JNK signaling in regulating spindle positioning during periods of tissue resizing and stem cell expansion becomes hijacked under conditions of stress and in the aging animal, limiting tissue homeostasis and shortening lifespan. It remains unclear how JNK is activated in ISCs during starvation / re-feeding, but insulin signaling has been implicated in promoting symmetric divisions during adaptive resizing of the *Drosophila* intestine (O’Brien et al., 2011). It will be interesting to test whether insulin signaling and JNK interact to regulate spindle positioning in ISCs, as elevated insulin signaling activity may also contribute to the age-related chronic activation of JNK. The age-related bias towards planar spindle orientations is reminiscent of the changes in spindle orientation of germline stem cells in old male flies (Cheng et al., 2008), and it is striking that restoring oblique spindle orientation in aged ISCs by increasing Kif1A expression is sufficient to improve intestinal physiology and extend lifespan. This indicates that the increases in ISC proliferation, which in the aging intestinal epithelium are a consequence of increased pro-inflammatory signaling (Li and Jasper, 2016), are not *per se* deleterious to tissue function, but that the de-regulation of cell specification events caused by JNK-induced changes in spindle orientation bias are more significant causes for epithelial dysfunction. This interpretation is supported by previous work in which it was shown that reducing Dl expression is sufficient to promote epithelial homeostasis in aging flies without restoring lower ISC proliferation rates (Biteau et al., 2008). Understanding the exact mechanisms and consequences of ISC spindle positioning is thus evidently critical to identify new intervention strategies to allay age-related dysfunction in barrier epithelia. Such interventions are likely to have significant clinical relevance, as barrier epithelia in mammals regenerate and age through mechanisms that are very similar to the *Drosophila* intestinal epithelium (Guo et al., 2014; Haller et al., 2017; Li and Jasper, 2016; Rock et al., 2011; Thevaranjan et al., 2017).

## Materials and Methods

### *Drosophila* stocks, husbandry, and treatments

The following UAS lines were obtained from the Bloomington *Drosophila* Stock Center: *w^1118^, bsk* DN (6409), *eb1-GFP* (35512), *luciferase* RNAi (31603), *GFP* (5431), *kif1a* FL (24787), *kif1a* RNAi (43264), *mud* RNAi (35044), *pins* RNAi (29310), *puc* RNAi (34392), and *wdr62* RNAi (53242). The following lines were gifts: *GFP-Mud* (Dr. Yohanns Bellïache), *hep* FL (Dr. Marek Mlodzik), *esg::gal4; Su*(*H*)*::Gal80, tub::Gal80^ts^* (Wang et al., 2014), *hs::FLP; FRT40, UAS::CD8-GFP* (Dr. Lucy O’Brien), *UAS::CD2-mir, UAS::CD2-RFP* (Dr. Lucy O’Brien), *hs::FLP; FRT40, tub::Gal80; tub::Gal4, UAS::GFP* (Dr. Benjamin Ohlstein), *FRT* UAS::GFP-mir *UAS-wdr62::mDendra2* (Dr. Clemens Cabernard), and *wdr62*^Δ3–9^ (Dr. Clemens Cabernard). The genotype of flies in each figure is detailed in Table S1.

Flies were maintained at 25°C and 65% humidity, on a 12-hour light/dark cycle. As needed, flies were shifted to 29°C for 4 days to induce genetic (TARGET) expression. Standard fly food was prepared with the following recipe: 1 l distilled water, 7.3 ml of EtOH, 13 g agar, 22g molasses, 65g malt extract, 18g brewer’s yeast, 80g corn flour, 10g soy flour, 6.2ml propionic acid, and 2g methyl-p-benzoate. For Geneswitch experiments, 200μM RU486 was additionally added. For Smurf experiments, 500mg/ml of Blue dye no. 1 (Alfa Chem) was additionally added. For lifespan and Smurf experiments, 75–125 female flies were housed together per bottle and flipped three times a week. Dead/Smurf flies were counted visually every other day.

For fast and refeed experiments, flies were fasted in 1% sucrose for one week, before returning to standard fly food for 3–7 days as noted. For Paraquat treatment, flies were starved for two hours before fed 5mM Paraquat in 5% sucrose on a vial containing Whatman filter paper for 20–24 hours, and intestines subsequently dissected. For Ecc15 infection, Ecc15 was cultured in 15ml of LB medium for 16 hours at 30° The culture was pelleted, resuspended in 5% sucrose, and fed to two-hour starved flies for 16–20 hours. Intestines were subsequently dissected. For activation of Twinspot and MARCM clones, flies were incubated at 37° for 45 minutes. Flies were Paraquat-treated or Ecc15-infected as described above just prior to heat shock and Twinspot analsysis was performed two days post heat shock. For MARCM clones, spindle orientation and clone size was quantified seven days post heat shock with Paraquat treatment performed a day before analyses.

### Live imaging of *Drosophila* intestine

Intestines from adult female flies were dissected in culture media containing 2mM CaCl_2_, 5mM KCl, 5mM HEPES, 8.2mM MgCl_2_, 108mM NaCl, 4mM NaHCO_3_, NaH_2_PO_4_ 10mM sucrose, 5mM trehalose, and 2% fetal bovine serum (Adult Hemolymph-like Saline, AHLS). Intestines were transferred to a 35mm glass bottom dish (MatTek, P35G-1.5-14-C), embedded in 4% low melting agarose (in AHLS), and submerged in AHLS. Intestines were imaged at intervals of 10 min for two hours on a Yokogawa CSUW1/Zeiss 3i Marianas spinning disk confocal microscopy system.

### Immunostaining of *Drosophila* intestine

Intestines from adult female flies were dissected in PBS (Phosphate-buffered saline, pH 7.4) and fixed for 25 minutes at room temperature in media containing 100mM glutamic acid, 25mM KCl, 20mM MgSO_4_, 4mM sodium phosphate, 1mM MgCl_2_, and 4% formaldehyde. Intestines were washed with PBS and stained in PBS, 0.3% Triton X-100 supplemented with 5% donkey serum. Intestines were incubated in primary antibody overnight at 4°C antibody, washed in PBS +0.01% Tween-20, and incubated in secondary antibodies, Phalloidin (1:400), and Hoechst stain (1:1000) for 2 hours at room temperature. For Delta staining, dissected intestines were incubated for ten minutes at room temperature in 100mM glutamic acid, 25mM KCl, 20mM MgSO_4_, 4mM sodium phosphate, 1mM MgCl_2_, and 4% formaldehyde, followed by 100% MeOH for five minutes at room temperature. Intestines were washed with 50% MeOH in PBS for two minutes, followed by PBS for five minutes. Immunostaining proceeded as described above.

Antibodies used in this study: Mouse monoclonal against Delta (gift from Dr. Matthew Rand), γ-tubulin (Sigma, T6557, RRID:AB_477584), phospho-JNK (Cell Signaling, 9255S, RRID:AB_2307321), and Wdr62 (gift from Dr. Clemens Cabernard); rabbit polyclonal against phospho-Histone H3 (EMD Millipore, 06-570, RRID:AB_310177); and rat monoclonal against α-tubulin (Bio-Rad MCA78G, RRID:AB_325005).

### FACs Sorting of ISCs and Quantitative Reverse Transcription PCR (RT-qPCR)

*w1118* or UAS-puc^RNAi^ flies were crossed to flies with nls-GFP expressing ISCs. Dissected intestines (100 per sample) were collected in 5% FBS in PBS on ice and treated with trypsin for 1 hr at 29°, followed by tituration to dissociate tissue. Cells were washed and resuspended in 5% FBS in PBS and GFP+ cells were sorted. RNA was extracted utilizing the RNeasy kit (Qiagen 74134) and converted to cDNA utilizing the iScript cDNA Synthesis kit (Bio-Rad 170-8890), according to manufacturers’ instructions. RT-qPCR was performed with cDNAs from ISCs from three populations (100 flies each) of controls and puc^RNAi^ expressing flies each on a QuantStudio 8 Flex system (ThermoFischer), using Taqman Probes (ThermoFischer): Kif1a/unc-104 (Dm01826374_g1) and Actin5c (Dm02361909_s1). For data analysis, C(t) values of Kif1a levels in linear scale were normalized to Actin5c.

### Microscopy and Image Analysis

Images were taken either with the Zeiss LSM 700 scanning confocal microscopy system or the Yokogawa CSU-W1/Zeiss 3i Marianas spinning disk confocal microscopy system. Intestines were imaged using 10x or 40x objective. 3-D reconstruction was performed using Icy (Quantitative Image Analysis Unit, Institut Pasteur, France). Images were analyzed using ImageJ (NIH, Bethesda, MD).

### Statistical Data

Each sample, ‘n’, represents the number of flies or cells as noted, from at least three independent experiments. Statistical analyses were performed with Prism (GraphPad Software, La Jolla, CA, USA). A two-sample Student’s t-test was used to compare means of two independent groups when the distribution of the data was normal. A one-way ANOVA with Tukey post-hoc test was used to determine statistical significance with multiple comparisons between three or more independent groups. A chi-squared test was used to test for statistical significance when comparing data according to a set hypothesis (i.e. percentage of spindles with phospho-JNK localization). A log-rank test was used to test for statistical significance in lifespan assays. Significance was accepted at the level of p < 0.05. No statistical methods were used to predetermine sample sizes, but our sample sizes are similar to those generally employed in the field. No randomization was used to collect data, but they were quantified blindly.

## Author Contributions

DJH and HJ conceived the project and wrote the manuscript; DJH performed experiments and analyzed data. All authors read and approved the final manuscript.

## Acknowledgement

We thank Drs. Yohanns Bellïache, Marek Mlodzik, Lucy O’Brien, Benjamin Ohlstein, and Clemens Cabernard for flies and reagents, and Nathaniel Troup for technical assistance. This project was supported by NIGMS (R01 GM117412) and NIA (R01 AG047497) to HJ, and NIA (F32AG053050) to DJH.

## Declaration of Interests

The authors declare no competing interests.

